# Targeting NAT10 Activates Tumor-Intrinsic Immunity and Suppresses Tumor Progression in Head and Neck Squamous Cell Carcinoma

**DOI:** 10.64898/2026.06.17.732930

**Authors:** Thurbu Tshering Lepcha, Nithya Paruchuri, Dawei Zhou, Guillaume N. Fiches, Jinshan He, K.A.S.N Shanaka, Yan Liu, Suha Eleya, Nischal Koirala, Jian Zhu, Darrion Mitchell, John W Zhao, Netty G. Santoso

**Author notes:** To whom correspondence should be addressed: Netty G. Santoso.

## Abstract

Head and neck squamous cell carcinoma (HNSCC) remain a major clinical challenge due to its high heterogeneity and limited therapeutic response, resulting in a 5-year overall survival rate of only ∼50%. Identifying molecular pathways that drive tumor progression while suppressing anti-tumor immunity is therefore critical for developing more effective therapies. N-acetyltransferase 10 (NAT10) is the only known enzyme responsible for catalyzing the RNA modification N4-acetylcytidine (ac4C) on rRNA, tRNA, and mRNA, and has been implicated in tumor progression in several cancers. In this study, we identify NAT10 as a key suppressor of tumor-intrinsic immune signaling in HNSCC. NAT10 expression was significantly elevated in tumor tissues and HNSCC cell lines and was associated with poor overall survival. Moreover, high-risk HPV, a major etiological factor in HNSCC, upregulated NAT10 protein expression through the viral oncoproteins E6 and E7. Functional inhibition of NAT10, either by genetic depletion or the small-molecule inhibitor Remodelin, activated tumor-intrinsic innate immune responses, as evidenced by increased IRF3 phosphorylation and induction of type I/II interferons and interferon-stimulated genes. Depletion of NAT10 was able to suppress tumorigenic phenotypes, including cell proliferation, migration, and colony formation in HNSCC cells. Importantly, activation of the STING signaling pathway using agonist cyclic di-GMP further amplified immune activation in NAT10-inhibited cancer cells. Together, our findings establish NAT10 as a previously unrecognized negative regulator of tumor-intrinsic immunity in HNSCC and support NAT10 targeting, particularly in combination with STING agonists, as a promising immunotherapeutic strategy.

## INTRODUCTION

Head and Neck Squamous Cell Carcinoma (HNSCC) arises from the mucosal epithelium of the oral cavity, pharynx, and larynx and represents the sixth most common cancer worldwide, with approximately 890,000 new cases and 450,000 deaths annually according to GLOBOCAN (1, 2, 3). Despite advances in diagnosis and treatment, the intrinsic heterogeneity and molecular complexity of HNSCC contribute to a persistently poor prognosis, with a 5-year overall survival (OS) rate of nearly 50%. Furthermore, the global burden of HNSCC continues to increase and is projected to rise by nearly 30% by 2030 (1). In Southeast Asia, the high prevalence of HNSCC is strongly associated with carcinogenic lifestyle factors such as cigarette smoking, smokeless tobacco use, excessive alcohol consumption, and betel nut chewing. Chronic exposure to these carcinogens promotes the accumulation of genetic and epigenetic alterations, including DNA damage, somatic mutations in key tumor suppressor genes such as TP53, deletions of CDKN2A, and amplification of oncogenes such as PIK3CA, ultimately driving malignant transformation in HNSCC (1).

In contrast to carcinogen-associated tumors, the incidence of Human Papillomavirus (HPV)–associated HNSCC has increased substantially in developed regions, including the United States and Europe (1, 2, 3). High-risk HPV subtypes, particularly HPV16 and HPV18, are predominantly associated with oropharyngeal squamous cell carcinomas. HPV-driven tumors exhibit distinct molecular and clinical characteristics compared with carcinogen-associated HNSCC, which more commonly arise in the oral cavity, hypopharynx, and larynx (1,2,3). Currently, the standard treatment for HNSCC involves surgical resection with or without chemotherapy and/or radiotherapy (4). Immunotherapy has also emerged as promising treatment option for patient with recurrent or metastatic disease. However, positive response rates are limited to a subset of patients and immune-related adverse effects can occur, underlying the need to better understand the regulation of tumor-intrinsic immune mechanism to develop strategies that can enhance immunotherapeutic efficacy.

Recently, we identified the histone methyltransferase PRDM6 as a novel epigenetic regulator that promotes tumor cell proliferation in HPV-positive HNSCC through an HPV E6/E7-dependent mechanism (5). Our previous work also demonstrated that PRDM6 suppresses tumor-intrinsic immune gene expression in HNSCC cells, representing a promising immunotherapeutic target for HPV-associated HNSCC (5). In the present study, we identify NAT10 (N-acetyltransferase 10), the only known enzyme responsible for catalyzing the N4-acetylcytidine (ac4C) modification of rRNA, tRNA, and mRNA, as an important epitranscriptomic regulator involved in immune signaling and tumorigenesis in HNSCC (6, 7). NAT10-mediated ac4C modification has been implicated in multiple cellular processes, including RNA stability, translation, and stress responses. Overall, aberrant NAT10 expression has been associated with oncogenic transformation, tumor cell proliferation, migration, and invasion in several cancers, including oral and laryngeal squamous cell carcinomas (8–11).

Epitranscriptomic regulation refers to post-transcriptional chemical modifications on RNA molecules that influence RNA stability, translation, and cellular signaling pathways. Increasing evidence indicates that these RNA modifications play important roles in immune regulation and tumor biology. For example, N6-methyladenosine (m6A), the most abundant internal RNA modification in eukaryotic mRNA, has been implicated in the regulation of immune responses and tumor progression in multiple cancers, including colorectal cancer, melanoma, and glioma (12). In addition, N1-methyladenosine (m1A) modifications have been detected in transcripts of interferon-stimulated genes, where they contribute to the regulation of type I interferon–mediated anti-tumor immunity (13). More recently, the RNA cytosine methyltransferases NSUN2, which catalyze the 5-methylcytosine (m5C) RNA modification, were shown to modulate activation of the cGAS–STING and the production of type I interferons, further highlighting the emerging role of epitranscriptomic regulation in innate immune signaling (14).

Type I interferons are central mediators of anti-tumor immunity that are produced by immune, stromal, and tumor cells and function primarily through the induction of interferon-stimulated genes (ISGs) (15). Activation of these genes modulates key cellular processes—including cell proliferation, survival, and angiogenesis—while also promoting anti-tumor immune responses mediated by T cells, natural killer cells, and dendritic cells (15, 16). In parallel, Interferon gamma (IFN-γ), the major type II interferon, further amplifies anti-tumor immunity by enhancing antigen presentation through upregulation of major histocompatibility complex molecules and promoting cytotoxic T-cell–mediated tumor cell elimination (17). A major upstream regulator of interferon signaling is the STING pathway, in which cytosolic DNA sensing, cGAS, activates STING and triggers recruitment of TBK1, leading to phosphorylation of IRF3 and transcription of interferons and ISGs (18, 19). Activation of this pathway has been shown to enhance anti-tumor immune responses and suppress tumor growth. In fact, pharmacologic STING agonists such as cyclic di-GMP have demonstrated promising anti-tumor efficacy in preclinical models, highlighting their potential as immunotherapeutic agents (20, 21, 22).

Here, we investigated the role of NAT10 in tumor-intrinsic immune regulation and tumorigenesis in Head and Neck Squamous Cell Carcinoma. Our findings demonstrate that NAT10 functions as an oncogene that is highly expressed in oral carcinoma cells. Inhibition of NAT10 significantly enhanced tumor-intrinsic interferon signaling, upregulated ISG expression, as well as inhibited tumor cell proliferation, migration, and clonogenic growth. We also evaluated the therapeutic potential of combining NAT10 inhibition with activation of STING using cyclic di-GMP. This combinatorial approach further amplified interferon responses and innate immune activation in HNSCC cells. These results identify NAT10 as a critical regulator of tumor-intrinsic immune signaling in HNSCC and targeting NAT10 may represent a promising strategy to enhance anti-tumor immunity.

## RESULTS

### N-Acetyltransferase 10 (NAT10) is up regulated in HNSCC cells

To investigate the role of NAT10 in head and neck cancer, we first compared its expression between cancerous and normal oral keratinocytes. RT-qPCR analysis revealed increased NAT10 transcript levels in both non-HPV and HPV-positive oral squamous cell carcinoma cell lines (SCC9, CAL27, and UM-SCC-47) compared with normal human oral keratinocytes (OKF4 and OKF6) (Figure 1A). Immunoblot analysis further demonstrated markedly elevated NAT10 protein levels in oral cancer cells relative to normal oral keratinocytes (Figure 1B). To further examine the contribution of high-risk HPV to NAT10 regulation, we evaluated NAT10 expression in HPV16-positive keratinocyte models. NAT10 protein levels were increased in HPV16-containing N/Tert-1 keratinocytes (N/Tert-1+HPV16) compared with HPV16-negative parental N/Tert-1 cells (Figure 1C). Similarly, ectopic expression of HPV16 E6, E7, or E6/E7 in 293T cells increased NAT10 protein abundance (Figure 1D). This regulation appeared to occur primarily at the post-transcriptional level, as NAT10 mRNA levels remained largely unchanged (Figure S1E, F).

**Figure 1.**
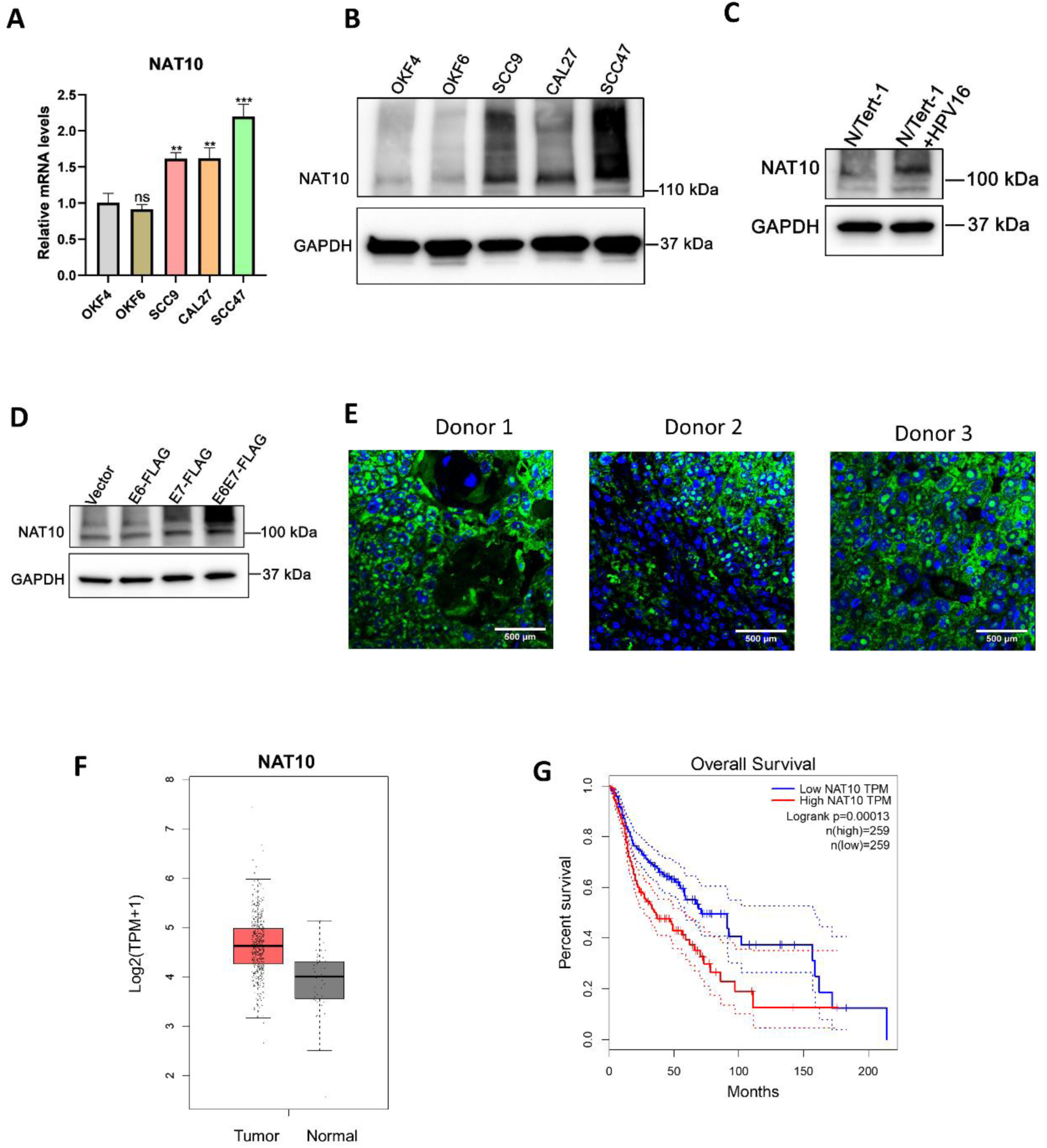
NAT10 expression is elevated in HNSCC. **(A, B**) NAT10 transcript and protein levels were analysed by RT-qPCR (A) or immunoblotting (B), respectively, in normal oral keratinocytes, and HNSCC cell lines. Equal protein loading was confirmed by reprobing the blots with a GAPDH antibody. **(C)** NAT10 protein expression in telomerase-immortalized normal human foreskin keratinocytes with or without HPV-16 genome (N/Tert-1+HPV16 and N/Tert-1, respectively) was analysed by immunoblotting. GAPDH was used as a loading control. **(D)** 293T cells were transfected with empty vector (vector) or with pLXSN vector expressing FLAG-tagged HPV16 E6, E7 and E6/E7, followed by immunoblotting analysis of NAT10 protein expression. GAPDH served as a loading control. **(E)** Representative immunofluorescence image showing NAT10 expression in HNSCC tissue microarray samples from three independent donors. Green indicates NAT10 staining, blue indicates Hoechst nuclear stain. Scale bar = 500 µm. **(F)** Analysis of TCGA dataset through GEPIA pipeline showing upregulation of NAT10 expression in HNSCC tumor compared with normal tissues. **(G)** Kaplan-Meier analysis demonstrating high NAT10 expression is associated with reduced overall survival in HNSCC patients. Statistical significance was determined using the log-rank test. For A, results represent mean ± S.D. (n = 3); **p < 0.01; ***p < 0.001; ns, nonsignificant. The blots in panels B, C and D are representative of results from three independent experiments.

We further confirmed high expression of NAT10 in pre-existing tumor tissues obtained from patients with HNSCC (Figure 1E). Consistent with these observations, analysis of the TCGA dataset demonstrated higher NAT10 mRNA levels in HNSCC tumor relative to adjacent normal tissues (Figure 1F). Moreover, Kaplan–Meier survival analysis revealed that high NAT10 expression significantly correlated with poorer overall survival in HNSCC patients (Figure 1G). Collectively, these findings confirmed NAT10 as a clinically relevant factor upregulated in both HPV-positive and HPV-negative HNSCC that supports its further mechanistic investigation in HNSCC.

### NAT10 suppresses the expression of innate immune response in HNSCC

Recent studies, including our own work, have implicated NAT10 in the regulation of innate immune signaling pathway (23). Innate immune responses—particularly those mediated through interferon signaling—play critical roles in anti-tumor immunity by promoting immune surveillance and restricting tumor growth (15, 16). One key pathway that drives interferon production is the cGAS–STING signaling pathway, which senses cytosolic DNA and activates downstream transcriptional programs that induce type I interferons and interferon-stimulated genes (24, 25). To investigate whether NAT10 modulates this pathway in HNSCC, we performed siRNA-mediated knockdown of NAT10 in SCC9 cells (Figure 2A and 2B) and CAL27 cells (Figure S1A) and examined activation of interferon signaling components. Depletion of NAT10 resulted in a pronounced increase in phosphorylated IRF3, the activated form of IRF3 that undergoes dimerization, nuclear translocation, and binding to interferon-stimulated response elements to drive transcription of interferon genes, while total IRF3 levels remained unchanged (Figure 2A). Consistent with activation of this pathway, NAT10 knockdown significantly increased the expression of both type I interferons (IFN-α and IFN-β) and the type II interferon, IFN-γ, in SCC9 cells (Figure 2B–D) and CAL27 cells (figure S1B-D). In addition, depletion of NAT10 led to elevated expression of interferon-stimulated genes, including ISG15 and IFIT1 (Figure 2F and 2G), further supporting activation of the interferon response program. To further determine whether NAT10 influences signaling upstream of STING, we treated control and NAT10-depleted SCC9 cells with the STING agonist cyclic di-GMP. Activation of STING with cyclic di-GMP resulted in an even greater induction of type I interferon expression in NAT10-depleted cells compared with control cells (Figure 2H), suggesting that loss of NAT10 sensitizes tumor cells to STING-mediated innate immune activation.

**Figure 2.**
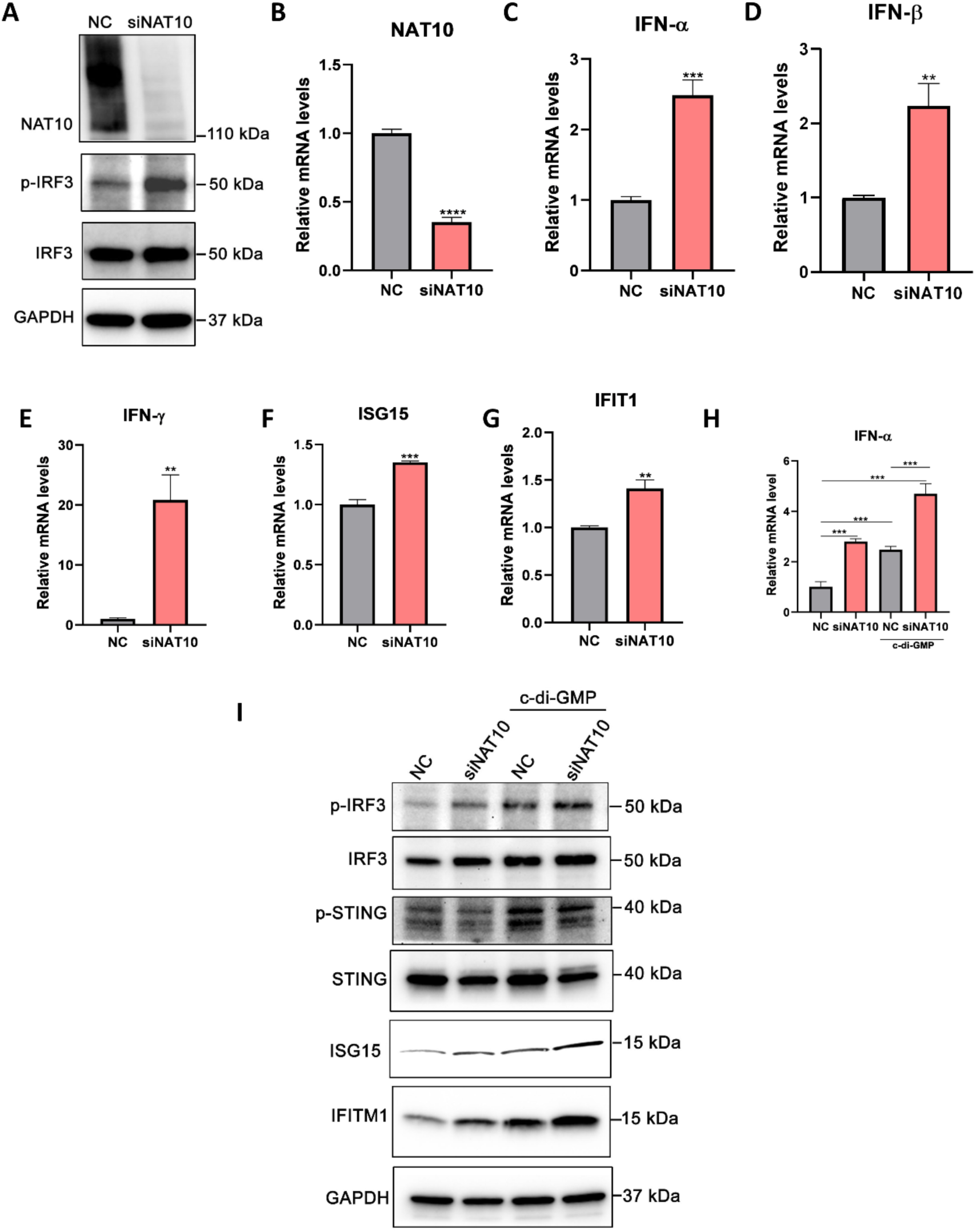
NAT10 regulates STING-mediated innate immune activation in HNSCC cells. **(A-G)** SCC9 cells were transfected with negative control siRNA (NC) or NAT10-targeting siRNA (siNAT10). Cell lysates were analyzed by immunoblotting with antibodies against NAT10, Phospho-IRF3, and IRF3. Equal loading was confirmed by GAPDH antibody (A). Total RNA was isolated, and transcript levels of IFN-α (B), IFN-β (C), IFN-γ (D), NAT10 (E), ISG15 (F) and IFITM1 (G) was quantitated by RT-qPCR. Results represent means ± S.D. (n = 3); ***p < 0.001; ns,nonsignificant. **(H, I)** SCC9 cells transfected with negative control siRNA (NC) or with siNAT10, were subsequently transfected with c-di-GMP, IFN-α transcript levels were quantified by RT-qPCR (H). Cell lysates were subjected to immunoblotting with NAT10, p-IRF3, IRF3, p-STING, STING, ISG15 and IFITM1 antibodies respectively (I). Equal loading for immunoblotting was confirmed by GAPDH expression. For panel H, data represents mean ± S.D. (n = 3); ***p < 0.001; ns, nonsignificant.

We further confirmed activation of interferon pathway through immunoblotting. Depletion of NAT10 resulted in increased levels of phosphorylated IRF3, phosphorylated STING, along with elevated expression of ISG15 and IFITM1, which were further enhanced by treatment with c-di-GMP (Figure 2I). Collectively, these findings indicate that NAT10 acts as a negative regulator of the cGAS–STING–IRF3 axis in HNSCC, thereby restraining interferon production and downstream antiviral-like immune responses. These results highlight NAT10 as an important modulator of innate immune signaling in HNSCC and suggest that targeting NAT10 may enhance anti-tumor immunity by potentiating cGAS–STING–dependent interferon responses.

### NAT10 inhibition by Remodelin triggers Type I and II interferon responses in HNSCC cells

Remodelin is a small-molecule inhibitor of NAT10 (48) that has been reported to exert promising antiviral and anticancer activities (26, 27). Type I and type II interferons (IFNs) are key mediators of host immune responses against pathogens and tumors and are primarily induced downstream of innate immune sensing pathways such as the cGAS–STING signaling pathway (19). Because our genetic studies showed that depletion of NAT10 upregulates type I and type II interferon signaling, we next sought to determine whether pharmacological inhibition of NAT10 using Remodelin similarly activates the cGAS–STING axis in HNSCC cells.

To address this, SCC9 oral cancer cells were treated with increasing concentrations of Remodelin (2.5, 5 and 10 μM). Remodelin treatment induced robust phosphorylation of IRF3 and increased expression of the interferon-stimulated protein ISG15 in a dose-dependent manner (Figure 3A), indicating activation of downstream innate immune signaling. We further confirmed that treatment with a higher dose of Remodelin (10uM) increased levels of phosphorylated IRF3 as well as the interferon-inducible protein IFITM1 (Figure 3B), while no significant cytotoxicity was observed under these treatment conditions (Figure S2A). Consistent with activation of interferon signaling, pharmacological inhibition of NAT10 also led to increased transcriptional levels of type I interferon (IFN-β) (Figure 3C), type II interferon (IFN-γ) (Figure 3D), and the interferon-stimulated gene ISG15 (Figure 3E) in SCC9 cells. Similar effects were observed in CAL27 cells as well (Figure S2B, S2C).

**Figure 3.**
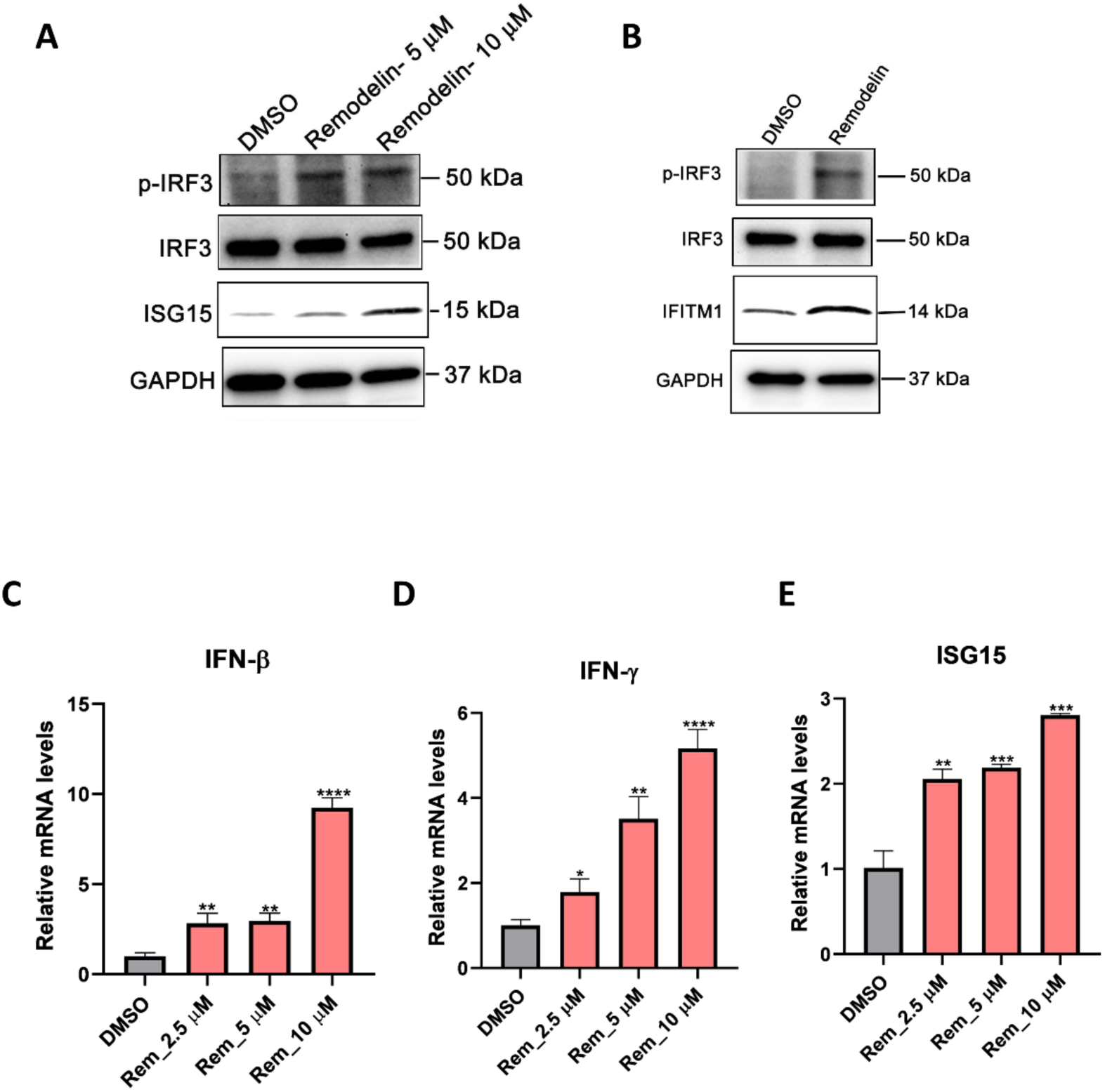
NAT10 regulates IRF3 activation in HNSCC cells. **(A)** SCC9 cells were treated with DMSO or Remodelin at different concentration followed by immunoblotting analysis using antibodies against p-IRF3, IRF3 and ISG15. **(B)** CAL27 cells were treated with DMSO or Remodelin (10 µM) followed by immunoblotting with p-IRF3, IRF3 and IFITM1 antibodies. GAPDH was used as a loading control. **(C, D, E)** Dose-dependent expression of IFN-β, IFN-γ and ISG15 transcripts were analysed by RT-qPCR in Remodelin-treated SCC9 cells. Results represent mean ± S.D. (n = 3); ***p < 0.001; ns, nonsignificant.

Taken together, these findings demonstrate that pharmacological targeting of NAT10 using Remodelin enhances activation of the cGAS–STING–IRF3 signaling cascade and amplifies downstream interferon responses in HNSCC cells. These results further support the role of NAT10 as a negative regulator of innate immune signaling and suggest that inhibition of NAT10 may represent a potential strategy to stimulate anti-tumor immunity in HNSCC.

### The combination of STING-agonist and Remodelin significantly enhances interferon responses

The STING signaling pathway is a critical regulator of interferon production and serves as an important bridge between innate and adaptive immune responses. To determine whether inhibition of NAT10 could cooperate with STING activation to enhance interferon signaling, we treated HNSCC cells with the NAT10 inhibitor Remodelin in combination with the STING agonist cyclic di-GMP and evaluated downstream immune signaling.

Our results showed that the immune response induced by Remodelin was further potentiated by the addition of cyclic di-GMP. Combined treatment led to robust activation of STING signaling, as evidenced by increased phosphorylation of IRF3 and enhanced expression of interferon-stimulated genes (ISGs) in HNSCC cells treated with Remodelin followed by cyclic di-GMP (Figure 4A). Consistent with this observation, co-treatment with NAT10 inhibition and STING activation synergistically elevated transcript levels of IFN-β (Figure 4B). Moreover, several downstream interferon-stimulated genes, such as ISG15, MX1, and OAS1, were significantly more upregulated following the combination treatment compared with Remodelin alone (Figure 4C–E).

**Figure 4.**
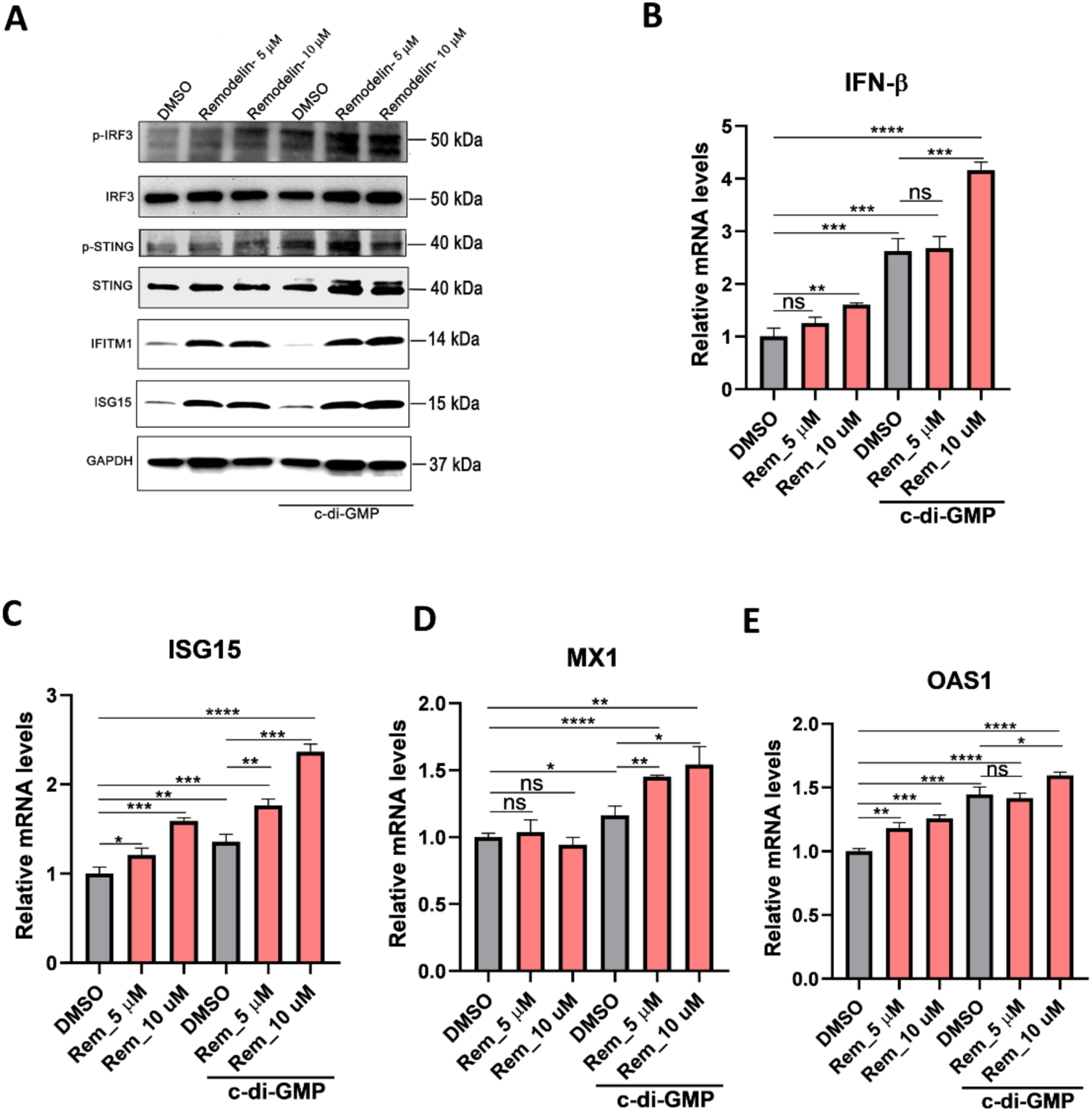
STING is activated in Remodelin-treated HNSCC cells in a cyclic-di-GMP-dependent manner. **(A-E)** SCC9 cells were treated with DMSO or Remodelin (10 µM), prior to transfection with cyclic-di-GMP. Cell lysates were analyzed by immunoblotting with p-IRF3, IRF3, p-STING, STING, IFITM1 and ISG15 antibodies (A). Transcript levels of IFN-β (B), ISG15 (C), MX1 (D) and OAS1 (E) were quantified by RT-qPCR. Equal protein loading was confirmed by reprobing of all blots with GAPDH antibody. Each blot is representative of three independent experiments. For B, C, D and E results represent mean ± S.D. (n = 3); **p < 0.01; ns, nonsignificant.

In addition, we observed a time-dependent induction of IFN-β, ISG15, and MX1 upon cyclic-di-GMP transfection in SCC9 cells (Figure S3A-C). Immunoblot data also showed that protein levels of STING and ISG15 activation peaked at 6 h and 24 h post-transfection with cyclic-di-GMP (Figure S3D). We further corroborated our results in CAL27 cells, where combined treatment with Remodelin and cyclic-di-GMP for 24 hours led to an increase of phospho-IRF3 expression, along with the subsequent induction of ISG15 and IFITM1 (Figure S3E).

Overall, these results demonstrate that pharmacological inhibition of NAT10 cooperates with STING activation to amplify interferon signaling and innate immune responses in HNSCC cells. These findings suggest that targeting NAT10 may sensitize tumor cells to STING-mediated immune activation and highlight a potential combinatorial therapeutic strategy to enhance anti-tumor immunity in HNSCC.

### NAT10 promotes the proliferation and migration of HNSCC cells in vitro

To determine the functional role of NAT10 in HNSCC tumorigenesis, we evaluated key malignant phenotypes including cell proliferation, migration, and clonogenic growth following pharmacological inhibition of NAT10. Treatment with the NAT10 inhibitor Remodelin significantly reduced cell proliferation in oral carcinoma cells SCC9 and CAL27 (Figure 5C, D), indicating that NAT10 activity contributes to cell growth.

**Figure 5.**
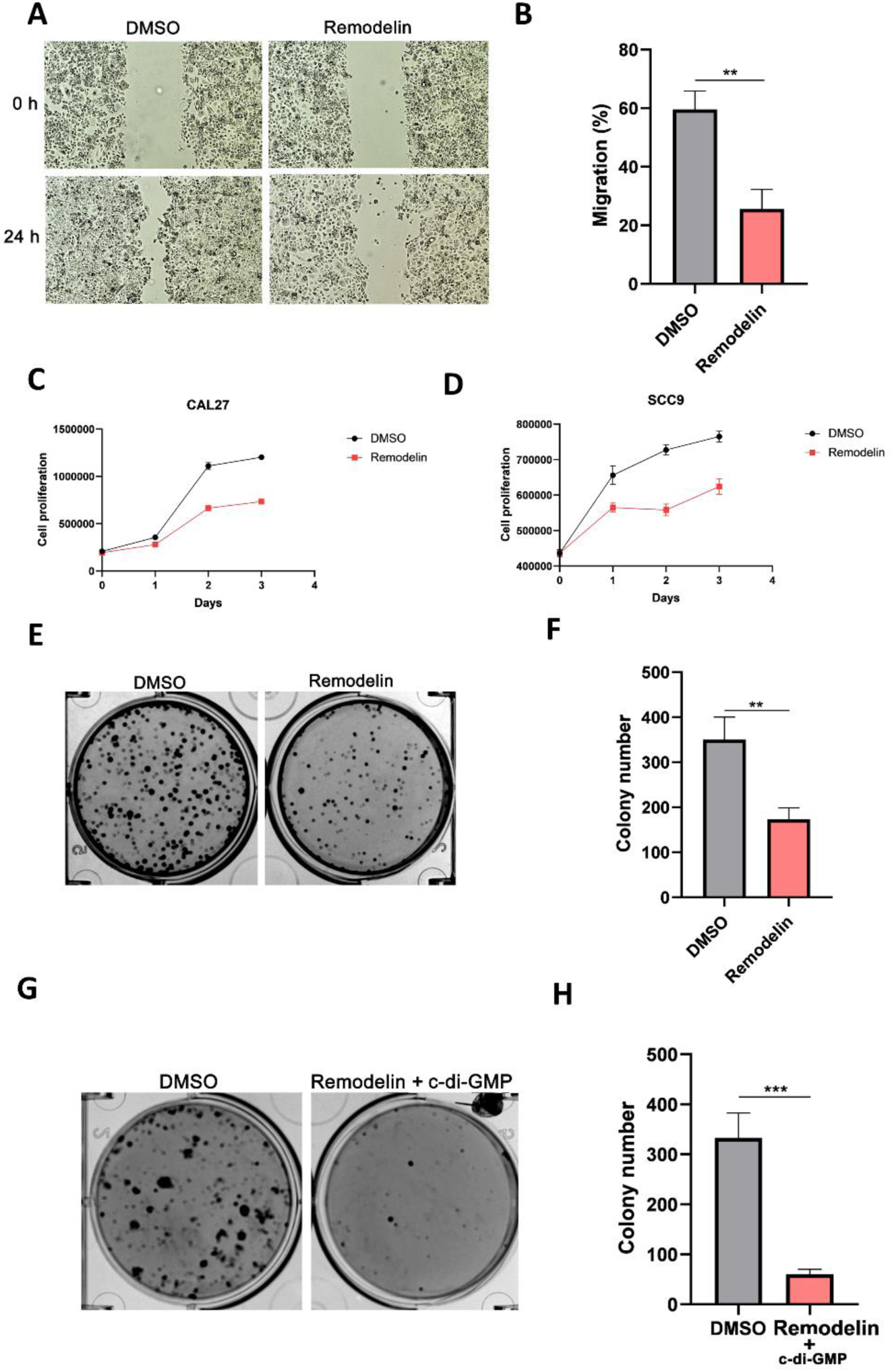
NAT10 regulates cell proliferation and migration in HNSCC cells. **(A, B)** CAL27 cells were treated with DMSO or Remodelin and subjected to a wound healing assay to assess cell migration capacity. **(C, D)** CAL27 and SCC9 cells were treated with either DMSO or Remodelin followed by the time-dependent analysis of cell proliferation. **(E-H)** SCC9 cells were treated with DMSO or Remodelin and colony formation ability was analysed (E, F) followed by cylic-di-GMP introduction (G, H). Results are representative of three independent experiments. DMSO = Dimethyl Sulfoxide.

We next assessed the impact of NAT10 inhibition on tumor cell motility using a wound-healing migration assay. Remodelin treatment markedly suppressed migration of SCC9 cells compared with control conditions (Figure 5A, B), suggesting that NAT10 may also contribute to the migratory capacity of HNSCC cells. Furthermore, inhibition of NAT10 significantly impaired clonogenic growth, as demonstrated by a marked reduction in colony formation by SCC9 cells (Figure 5E, F), a hallmark of tumorigenic potential. Similarly, we confirmed the reduction of cell survival and cell proliferation capacity with NAT10 inhibition in CAL27 cells as well (Figure S4A-D).

Interestingly, this anti-tumor phenotype was further enhanced when NAT10 inhibition was combined with activation of the cGAS–STING signaling pathway using a cGAS agonist (Figure 5G, H). The combination treatment resulted in a more pronounced suppression of clonogenic growth, consistent with enhanced activation of innate immune signaling and downstream interferon responses. Collectively, these findings demonstrate that targeting NAT10 suppresses multiple malignant properties of HNSCC cells—including proliferation, migration, and anchorage-independent growth—and suggest that these effects are at least partially mediated through modulation of interferon signaling downstream of the cGAS–STING pathway.

## DISCUSSION

NAT10 is a post-transcriptional gene regulator that catalyzes the addition of an acetyl group to the N4 position of cytidine, generating the RNA modification N4-acetylcytidine (ac4C) (28). Dysregulation of NAT10 has been reported in multiple malignancies, including hepatocellular carcinoma, gastric cancer, colorectal cancer, pancreatic cancer, breast cancer, multiple myeloma, and acute myeloid leukemia, where it contributes to tumor progression (29). In head and neck cancers, NAT10 is frequently overexpressed and promotes tumorigenesis in laryngeal squamous cell carcinoma by inhibiting pyroptosis and acetylating the 3′UTR of FOXM1 mRNA, thereby stabilizing this oncogenic transcription factor (30). NAT10 has also been shown to acetylate and stabilize mRNA encoding GLMP (glycosylated lysosomal membrane protein), leading to activation of the MAPK/ERK signaling pathway in HNSCC (31). In addition, NAT10 promotes cell proliferation, migration, and invasion in oral squamous cell carcinoma through ac4C-mediated acetylation of MMP1 mRNA (32).

Despite these findings, the functional role of NAT10 in immune regulation within head and neck squamous cell carcinoma remains poorly understood. Immunotherapy has recently emerged as a promising treatment strategy for patients with recurrent or metastatic HNSCC, in particular HPV-negative cases, however only a limited proportion of patients achieve durable responses (33). One possible contributor to this limited efficacy is the concurrent use of conventional therapies that may suppress or dampen anti-tumor immune responses (34). Consequently, ongoing clinical trials are exploring combination approaches incorporating neoadjuvant or adjuvant treatments to enhance therapeutic response rates and improve survival outcomes. Recent studies suggest that NAT10 may influence antitumor immunity and the tumor microenvironment, highlighting its potential as a target to improve cancer immunotherapy (35–37).

In this study, we investigated the role of tumor-intrinsic NAT10 in regulating immune responses in HNSCC. Consistent with previous reports, we observed that NAT10 expression is significantly elevated in tumor epithelial cells compared with normal oral epithelial cells (38, 39). Further, we linked the upregulation of NAT10 to infection of high-risk HPV. Importantly, our findings reveal a previously unrecognized link between NAT10 and tumor-intrinsic immune signaling. Suppression of NAT10 increased phosphorylation of IRF3 and enhanced expression of interferons, including IFN-α, IFN-β, and Interferon gamma. This activation of interferon signaling subsequently induced the expression of anti-tumor interferon-stimulated genes such as ISG15, IFITM1 and IFIT1 (23), suggesting that NAT10 functions as a negative regulator of innate immune signaling in HNSCC.

Pharmacological inhibition of NAT10 using the small-molecule inhibitor Remodelin activated both type I and type II interferon signaling in HNSCC tumor cells. Treatment of oral squamous cell carcinoma (OSCC) cells with Remodelin also impaired tumor cell proliferation and migration, supporting a functional role for NAT10 in promoting malignant phenotypes in HNSCC. Although the precise mechanisms underlying Remodelin activity in HNSCC require further investigation, previous studies have identified MYC as a key downstream target of NAT10, whose mRNA stability is enhanced through ac4C modification (27). Inhibition of NAT10 by Remodelin disrupts the MYC/CDK2/DNMT1 signaling axis and promotes accumulation of double-stranded RNA, which can trigger type I interferon responses and enhance tumor-specific CD8⁺ T-cell immunity (27). In addition to type I interferons, interferon gamma is known to exert anti-tumor effects by enhancing cytotoxic immune responses, inhibiting angiogenesis, and increasing expression of major histocompatibility complex molecules, which also facilitates immune-mediated tumor elimination (40–42).

Because IRF3 activation is a central component of the STING signaling pathway, we next examined whether pharmacologic activation of STING could further enhance immune responses induced by NAT10 inhibition. Loss or dysfunction of STING signaling is known to enable tumor cells to evade immune surveillance, whereas activation of STING can increase tumor immunogenicity and stimulate anti-tumor immune responses. Accordingly, treatment with the STING agonist, cyclic di-GMP, further potentiated interferon signaling in NAT10-inhibited HNSCC cells. Cyclic di-GMP is a bacterial cyclic dinucleotide that functions as a second messenger in bacteria but can also activate STING in mammalian cells, inducing robust type I interferon responses (43–47). Cyclic dinucleotides have therefore emerged as promising immunotherapeutic agents and vaccine adjuvants, with demonstrated ability to enhance anti-tumor immunity in multiple cancer models, including melanoma, where cyclic di-GMP promotes interferon production and activation of natural killer cells (47). Consistent with these immunostimulatory effects, combined treatment with the NAT10 inhibitor Remodelin and cyclic di-GMP further enhanced tumor-intrinsic interferon responses in HNSCC cells compared with either treatment alone. These findings suggest that NAT10 functions as a suppressor of tumor-intrinsic immune signaling and that its inhibition may sensitize HNSCC cells to STING-mediated immune activation. Thus, combining NAT10 inhibition with STING agonists may represent a promising strategy to enhance anti-tumor immunity and potentially improve the efficacy of existing immunotherapies, including immune checkpoint blockade. This approach may be particularly relevant to HPV-negative HNSCC, which is associated with poorer clinical outcomes and has limited therapeutic options, especially in the recurrent or metastatic setting.

Collectively, our findings demonstrate that NAT10 is dysregulated in both HPV-positive and HPV-negative HNSCC and contributes to the suppression of tumor-intrinsic immune responses. Although the precise molecular mechanisms through which NAT10 inhibits IRF3 activation and downstream type I and type II interferon signaling remain to be elucidated, its established role as the primary writer of RNA N4-acetylcytidine (ac4C) modifications suggests that NAT10 may promote the stability and/or translation of specific target mRNAs encoding negative regulators of innate immune signaling. Further studies are warranted to define the molecular mechanisms underlying NAT10-mediated immune suppression, identify the critical ac4C-modified target transcripts, and evaluate the therapeutic efficacy and safety of NAT10-targeting strategies alone or in combination with STING agonists and existing immunotherapies. These studies should employ diverse pre-clinical models of HNSCC, with particular emphasis on the HPV-negative disease, where the development of more effective therapeutic approaches remains an important unmet clinical need.

## METHODS

### Reagents

The following antibodies were used in the study: NAT10 (Cat # 13365-1-AP, Proteintech), ISG15 (Cat # sc-166755, Santa Cruz Biotechnology), IFITM1 (Cat # 60074-1-lg,Proteintech), IRF3 (Cat # 4302S, Cell Signaling), Phospho-IRF3 (Ser386) (Cat # 37829S, Cell Signaling), GAPDH (Cat # sc-47724, Santa Cruz Biotechnology), and horseradish peroxidase-tagged secondary antibodies HRP-conjugated anti-mouse IgG antibody (Cat. # 7076S), and HRP-conjugated anti-rabbit IgG antibody (Cat. # 7074S), were obtained from Cell Signalling Technology (CST). Alexa 488-conjugated anti-rabbit secondary antibody was from Invitrogen (Molecular Probes). Remodelin small molecular NAT10 inhibitor was purchased from MedChemExpress (Cat # HY16706A,). STING agonist c-di-GMP was purchased from TOCRIS biotechne (Cat# 5900).

### Cell culture

Human squamous carcinoma cell lines, CAL 27 (ATCC CRL-2095) and SCC9 (CRL-1629), were purchased from the American Type Culture Collection (ATCC). UM-SCC-47 (Cat. # SCC071) cells were kindly provided by Dr. Darrion Mitchell from Ohio State University. CAL27 and UM-SCC-47 (SCC47) cells were maintained in Dulbecco’s Modified Eagle Medium (DMEM; Gibco) supplemented with 10% heat-inactivated fetal bovine serum (FBS) and 1× penicillin–streptomycin at 37 °C in a humidified incubator with 5% CO₂. SCC9 cells were cultured in a 1:1 mixture of DMEM and Ham’s F12 medium containing 1.2 g/L sodium bicarbonate, 2.5 mM L-glutamine, 15 mM HEPES, and 0.5 mM sodium pyruvate, supplemented with 400 ng/mL hydrocortisone and 10% heat-inactivated FBS.

Normal human keratinocytes, keratinocytes immortalized by transfection either to express TERT (50,51). OKF4/Tert-1 (50) and OKF6/Tert2 cells (51) were kindly shared by Dr. James G. Rheinwald and were cultured in Keratinocyte SFM Kit (Cat. #17005042) and in Defined Keratinocyte SFM (Cat. #10744019) medium, respectively. All cells were maintained at 37 °C in a humidified atmosphere containing 5% CO₂. N/Tert-1 are Normal foreskin Keratinocytes immortalised with hTert, cultured in complete Keratinocyte serum free media (Invitrogen; 37010-022) with hygromycin. N/Tert-1+Xw14 - a cell line derived from the N/Tert-1, expressing HPV16 full genome and cultured in complete kSFM. Both N/Tert-1 and N/Tert-1+HPV16 cells (52) were kindly shared by Dr. Iain Morgan from Virginia Commonwealth University.

### Cell transfection

Reverse transfection of NAT10 siRNA (Cat. #S30492, Life Technologies) or non-targeting control siRNA (Cat. #AM4641) was performed using Lipofectamine RNAiMAX reagent (Cat. #13778030, Invitrogen) according to the manufacturer’s instructions, and cells were incubated for 72 h. The STING agonist cyclic di-GMP (c-di-GMP) was transfected using FuGENE HD transfection reagent (Cat. #E2691, Promega).

### Western blotting

Following transfection or treatment, cells were washed with cold PBS and lysed in 1× RIPA buffer (Cat. #20-188, Millipore) supplemented with protease inhibitor cocktail (Cat. #A32965, Thermo Scientific) for 20 min on ice. Protein concentrations were determined using a BCA protein assay kit (Cat. #23225, Thermo Scientific). Lysates were centrifuged at 10,000 × g for 10 min at 4 °C, and supernatants were collected. Samples were mixed with SDS loading buffer containing 5% β-mercaptoethanol and resolved on Mini-PROTEAN TGX gels (Bio-Rad). Proteins were transferred to PVDF membranes, blocked with 5% nonfat dry milk, and incubated overnight at 4 °C with primary antibodies, followed by HRP-conjugated secondary antibodies. Immunoreactive bands were visualized using Clarity Max ECL substrate (Cat. #1705062, Bio-Rad).

### Quantitative RT-PCR

Total RNA was extracted from HNSCC cells using the NucleoSpin RNA extraction kit (Cat. #740955.250, MACHEREY-NAGEL) according to the manufacturer’s instructions. cDNA synthesis was performed using the iScript cDNA Synthesis Kit (Cat. #1708890, Bio-Rad). Quantitative real-time PCR (RT-qPCR) was carried out using iTaq Universal SYBR Green Supermix (Cat. #1727125, Bio-Rad) on a Bio-Rad CFX Connect system. The thermal cycling conditions were as follow: an initial denaturation at 95 °C for 10 min, followed by 50 cycles of 95 °C for 15 s and 60 °C for 1 min. GAPDH was used as the internal normalization control. Relative gene expression was calculated using the comparative ΔΔCt method, and fold changes were determined as 2−ΔΔCt. Primer sequences are listed in Table S1.

### Protein immunofluorescence

Paraffin-embedded HNSCC tissue sections were baked at 65 °C for 30 min, followed by deparaffinization in xylene and graded ethanol washes before rehydration in distilled water. Antigen retrieval was performed using a 2100 Retriever system (Electron Microscopy Sciences, Cat. #62700-10) with Tris-EDTA antigen retrieval buffer (pH 9.0; Abcam, Cat. #ab93684) for 2 hrs to complete the cycle and then cool down. To block nonspecific binding, sections were incubated with 10% normal goat serum (NGS) in PBST for 2 h at room temperature. Slides were then incubated overnight at 4 °C with rabbit anti-NAT10 primary antibody diluted in 5% NGS/PBST. After washing, sections were incubated with Alexa Fluor 488-conjugated anti-rabbit secondary antibody for 1 h at room temperature. Nuclei were counterstained with Hoechst 33342 for 15 min. Coverslips were mounted using ProLong Glass Antifade Mountant (Invitrogen, Cat. #P36982) and allowed to cure overnight in the dark. Images were acquired using a ZEISS LSM 700 confocal microscope and processed with ZEN imaging software.

### Wound healing assay

HNSCC cell migration was assessed using a wound healing assay. Briefly, 5 × 10^5^ cells were seeded into 6-well plates and allowed to reach ∼95% confluency prior to treatment with Remodelin. A scratch wound was generated using a 200 μL pipette tip, and detached cells were removed by washing with serum-free medium. Images were captured immediately after wounding (0 h) under a phase-contrast microscope to measure the initial wound area. Cells were then incubated with either DMSO control or Remodelin at 37 °C with 5% CO₂ for 24 h to allow migration into the wound area, followed by imaging at the endpoint. Migration was quantified using the following formula:

[(wound distance at 0 h) - (wound distance at 24 h)] / [wound distance at 0 h] (49).

### Cell viability assay

Cell viability and proliferation were measured using the ATP-based CellTiter-Glo assay (Cat. #G7571, Promega) according to the manufacturer’s instructions. Briefly, HNSCC cells were seeded in 96-well plates and treated with Remodelin for the indicated time points. CellTiter-Glo reagent was then added directly to the culture medium at a 1:1 ratio in opaque 96-well plates. Luminescence was measured using a Cytation 5 plate reader.

### Colony formation Assay

HNSCC cells treated with Remodelin or DMSO were seeded into 6-well plates at a density of 1,000 cells per well and cultured for 14 days at 37 °C with 5% CO₂. Colonies were fixed with 4% paraformaldehyde and stained with 0.2% crystal violet for 30 min. Colony images were captured, and colony numbers were manually counted from three independent experiments.

### Statistical analysis

Statistical comparisons between two groups were performed using Student’s *t*-test. A *p* value < 0.05 was considered statistically significant.

## ACKNOWLEDGEMENT

We thank Dr. James Rheinwald for providing OKF4/Tert-1 and OKF6/Tert-2 cells. We also acknowledge Dr. Iain Morgan for sharing N/Tert-1 and N/Tert-1+HPV-16 cells.

## DATA AVAILABILITY

All data generated and/or analysed during this study are included in this published article and its supplementary materials. Any additional datasets supporting this study are available from the corresponding author upon reasonable request.

---

**Supplementary Figure 1.**
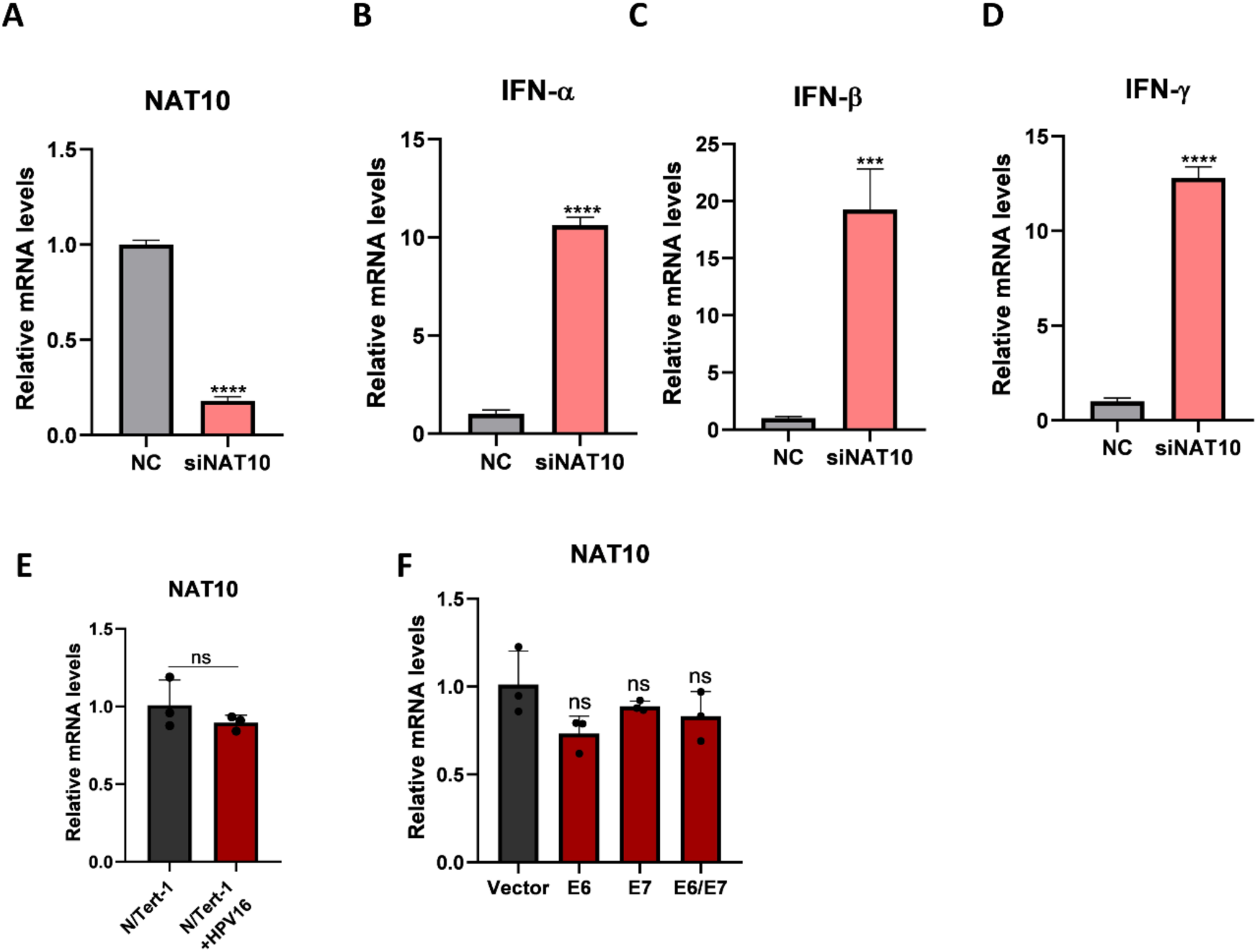
Knockdown of NAT10 increases interferon expression in HNSCC cells. **(A-D)** CAL27 cells were transfected with either a negative control (NC) or siNAT10. Following cell lysis, RNA was isolated, and transcript levels of NAT10, IFN-α, IFN-β, and IFN-γ were analysed by RT-qPCR (A, B, C and D). **(E)** Total RNA was extracted from telomerase-immortalized human foreskin keratinocytes with or without the HPV16 genome (N/Tert-1+HPV16 and N/Tert-1, respectively), followed by RT-qPCR analysis of NAT10 transcripts normalized to GAPDH. **(F)** 293T cells were transfected with empty vector (vector) or with pLXSN vector expressing FLAG-tagged HPV16 E6, E7 and E6/E7, followed by RT-qPCR analysis of NAT10 mRNA expression. Results represent mean ± S.D. (n = 3); ***p < 0.001; ns, nonsignificant.

**Supplementary Figure 2.**
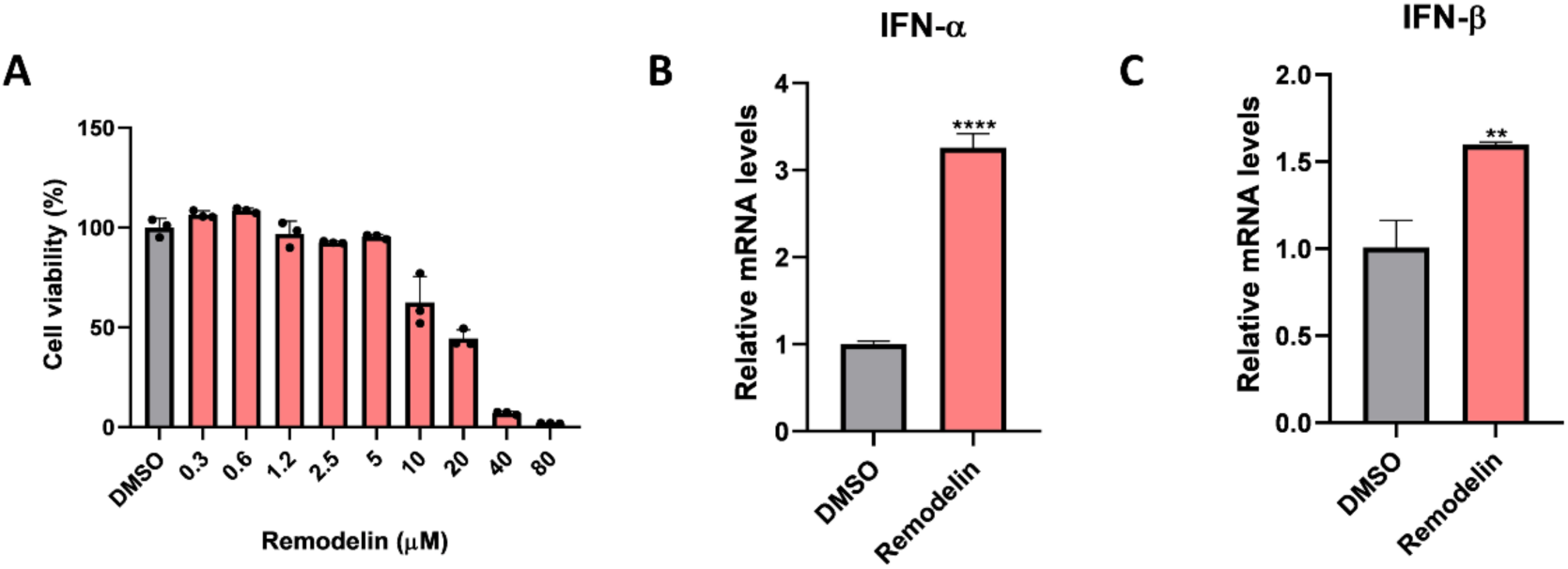
Inhibition of NAT10 using Remodelin in HNSCC cells. **(A)** SCC9 cells were treated with different doses of Remodelin for 48 hr followed by a cell viability assay using Cell Titer-Glo Luminescent Cell Viability Assay. **(B, C)** Transcript levels of IFN-α and IFN-β were quantified by RT-qPCR following Remodelin treatment in CAL27 cells. Results represent mean ± S.D. (n = 3); ***p < 0.001; ns, nonsignificant.

**Supplementary Figure 3.**
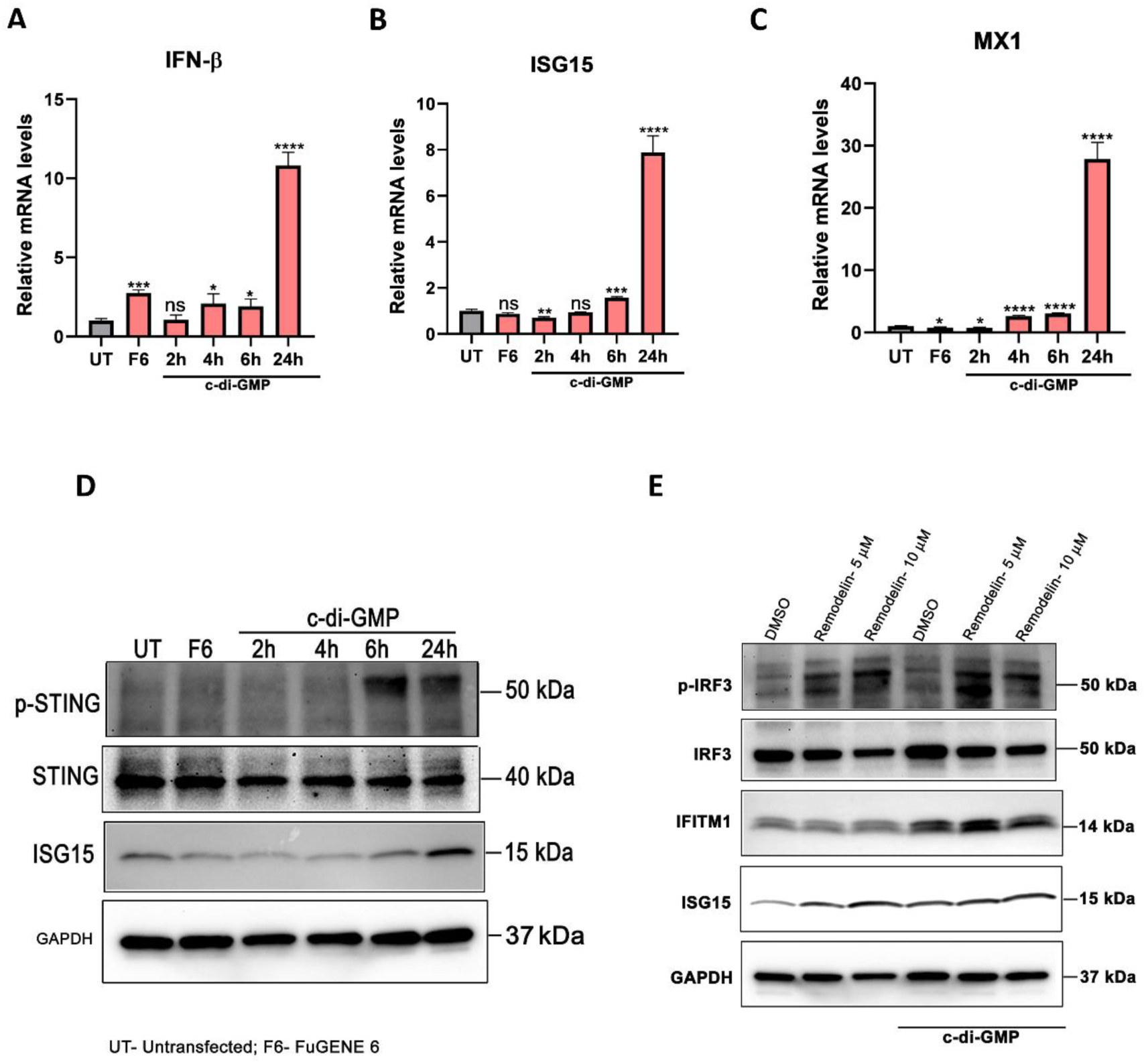
Time-dependent induction of interferons and ISGs in HNSCC cells following cyclic-di-GMP transfection. **(A-C)** Time-dependent expression of IFN-β (A), ISG15 (B) and MX1 (C) were analysed by RT-qPCR in cylic-di-GMP-transfected SCC9 cells. **(D)** Expression of p-STING and ISG15 were analysed in SCC9 cells by immunoblotting of cell lysates with anti-p-STING, STING and ISG15 antibodies. **(E)** CAL27 cells were treated with either DMSO or Remodelin, followed by cyclic-di-GMP transfection for 24 h. Cells were lysed, and protein levels of p-IRF3, IRF3, IFITM1 and ISG15 were analysed using immunoblotting with respective antibodies (E). Equal protein loading was confirmed by reprobing of all blots with GAPDH antibody. For A, B and C, results represent mean ± S.D. (n = 3); ***p < 0.001; ns, nonsignificant. The blots in panels D and E are representative of the results obtained in three independent experiments. UT, Untransfected; F6, FuGENE 6.

**Supplementary Figure 4.**
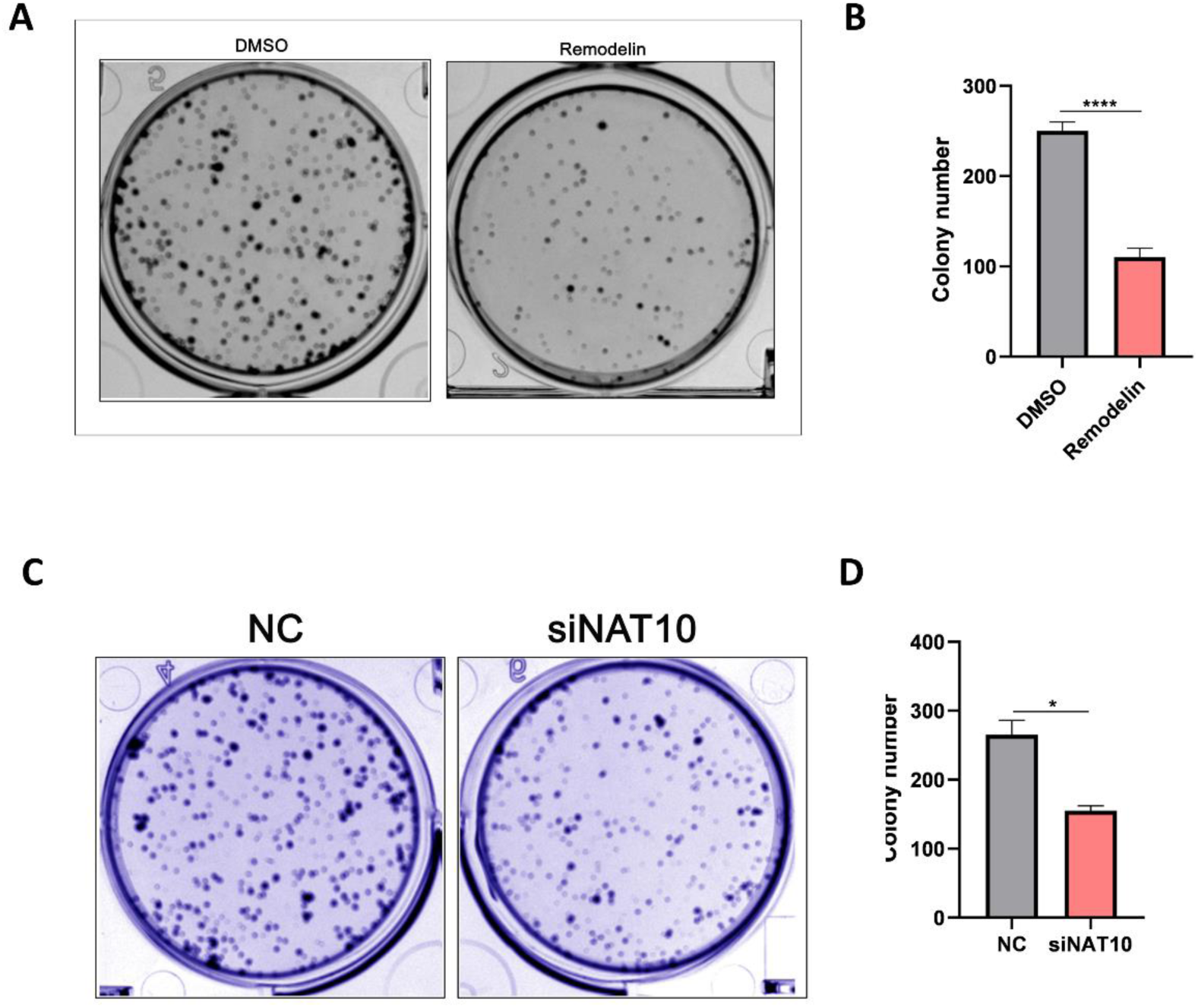
Knockdown or inhibition of NAT10 reduces cell proliferation. **(A-D)** CAL27 cells were transfected with DMSO or Remodelin (A, B), or negative control siRNA (NC) or siNAT10 (C, D), and subsequently subjected to colony formation assay. Representative images and quantifications from three independent experiments are shown.

